# BTFBS: binding-prediction of bacterial transcription factors and binding sites based on deep learning

**DOI:** 10.1101/2024.09.19.613986

**Authors:** Bingbing Jin, Song Liang, Xiaoqian Liu, Rui Zhang, Yun Zhu, Yuanyuan Chen, Guangjin Liu, Tao Yang

## Abstract

**Background:** The binding of transcription factors (TFs) to TF-binding sites plays a vital role in the process of regulating gene expression and evolution. With the development of machine learning and deep learning, some successes have been achieved in predicting transcription factors and binding sites. Then a natural question arises: for a given transcription factor and a binding site, do they bind? This is the main motivation of this work.

**Results:** In this paper, we develop a model BTFBS, which predicts whether the bacterial transcription factors and binding sites combine or not. The model takes both the amino acid sequences of bacterial transcription factors and the nucleotide sequences of binding sites as inputs, and extracts features through convolutional neural network and MultiheadAttention.

For the model inputs, we use two negative sample sampling methods: RS and EE. On the test dataset of RS, the accuracy, sensitivity, specificity, F1-score and MCC of BTFBS are 0.91446, 0.89746, 0.93134, 0.91264 and 0.82946, respectively. And on the test dataset of EE, the accuracy, sensitivity, specificity, F1-score and MCC of BTFBS are 0.87868, 0.89354, 0.86394, 0.87996 and 0.75796, respectively. Meanwhile, our findings indicate that the optimal approach for obtaining negative samples in the context of bacterial research is to utilize the whole genome sequences of the corresponding bacteria, as opposed to the shuffling method.

**Conclusions:** The above results on the test dataset have shown that the proposed BTFBS model has a good performance in predicting the combination of bacterial transcription factors and their binding sites and provides an experimental guide. BTFBS is publicly available at https://github.com/Vceternal/BTFBS.

## Background

In the network of prokaryotic transcriptional regulation, one of the most common forms is a transcription factor (TF), which has a single polypeptide containing both a sensory domain and a DNA-binding helix-turn-helix domain that directly binds DNA located at the promoter region of the gene (the “Switch” of gene transcription), thus affecting the binding of RNA polymerase to the promoter and in turn controlling gene transcription [1]. Although transcription factors binding data obtained by experimental methods is of high accuracy, it is time-consuming, laborious and expensive. Therefore, we hope to use computational methods to assist experimental screening and reduce research costs.

With the continuous development of machine learning and deep learning, more and more researchers use them to predict the binding sites of transcription factors. For example, DeepBind was the first deep learning method to predict protein-DNA-binding sites using convolutional neural networks [2]. Dilated [3] was a deep learning approach to model long-distance sequences by using dilated convolutional neural networks. KEGRU[4] combined the Bidirectional Gated Recurrent Unit (GRU) network with k-mer embedding to identify transcription factor binding sites. DanQ, a hybrid convolution and recurrent network framework, was a valuable resource for exploring the utility of non-coding DNA [5]. MAResNet was a deep learning method that combines the bottom-up and top-down attention mechanisms and a state-of-the-art feed-forward network (ResNet), which was bulit by stacking attention modules that generate attention-aware features [6]. Machine learning and deep learning have obtained significant achievements in the prediction of binding sites.

There are also approaches focusing on the prediction of transcription factors. DeepTFactor employed convolutional neural network to extract protein features for the purpose of predicting transcription factors [7]. By constructing a bacterial transcription factors database, Oliveira Monteiro et al. trained the deep learning model PredicTF, which provided the first pipeline for predicting and annotating TF within complex microbial communities [8]. However, these models only considered the prediction and classification of bacterial transcription factors, and did not take their binding sites into account.

In short, there are numerous methods to predict transcription factors and binding sites. Then there is a natural question: for e.g. bacteria, given a transcription factor and a binding site, how to determine whether they bind or not? This paper constructs a prediction model using deep learning to give an answer to this question.

As we know, the core idea of deep learning is to extract features and build models by training large-scale data. Data is the cornerstone of deep learning, containing rich information and rules, and is the original material for model learning and optimization. Data of high quality can provide more accurate and comprehensive information, thereby enhancing the performance and generalization ability of the model. For the prediction of transcription factor binding sites, positive samples can be obtained from experimental papers and from some databases, which unfortunately do not provide negative samples. Therefore, the selection of negative samples plays a crucial role in the prediction performance of the model. Some researchers use the shuffling method to obtain negative samples, such as DeepBind [2], MAResNet [6], deepRAM [9]. But the principle of this method is to shuffle the positive sequences while matching the dinucleotide composition [9] and the negative samples are probably not in the original whole genome sequences. To overcome this limitation, we use two negative sample sampling methods to select the negative samples from the whole genome sequences of the corresponding bacteria.

In general, based on convolutional neural network (CNN), ordinal positional encoding [10] and MultiheadAttention, we construct a deep learning model named BTFBS .

## Methods

### Datasets

At present, prokaryotes lack a comprehensive database of transcriptional regulation as eukaryotes. CollecTF [11] is an experimentally verified database of transcription factor-binding sites in bacteria. As CollecTF provides the correspondence between transcription factor (TF) accession numbers and the binding sites, it is necessary to retrieve the amino acid sequences based on the TF accession numbers. It is regrettable that CollecTF has only provided data up to 2015. PRODORIC is one of the largest collections of prokaryotic transcription factor binding sites from a multitude of bacterial sources [12]. Our data from 2016 to 2021 is obtained from this database. As a result, 5159 non-redundant positive sequences are obtained. Approximately 10% of the sequences are randomly selected from the datasets as the independent dataset, while the remaining 90% of the sequences are used as the training and test datasets.

For negative samples, we adopt two methods. The first method (RS) is to randomly select negative samples from the whole genome sequences of the bacteria to which the binding sites belong [13]. Given that the binding regions of bacterial transcription factors are predominantly located within non-coding regions, we only randomly intercept the non-coding regions from the whole genome sequences of the corresponding bacteria. Difflib (a flexible python library) is employed for the comparison of sequence similarity between the intercepted sequences and the binding sites, thus avoiding the inadvertent capture of the latter. The sequence is excluded if the similarity score between the two sequences exceeds 90%. Subsequently, 27,236 non-redundant sequences are obtained, and 19% of the data are randomly selected as negative samples, matching the number of positive samples. Approximately 10% of the sequences are randomly selected as the independent dataset, while the remaining 90% are used as the training and test datasets. Figure 1A illustrates the process of RS.

**Figure1.**
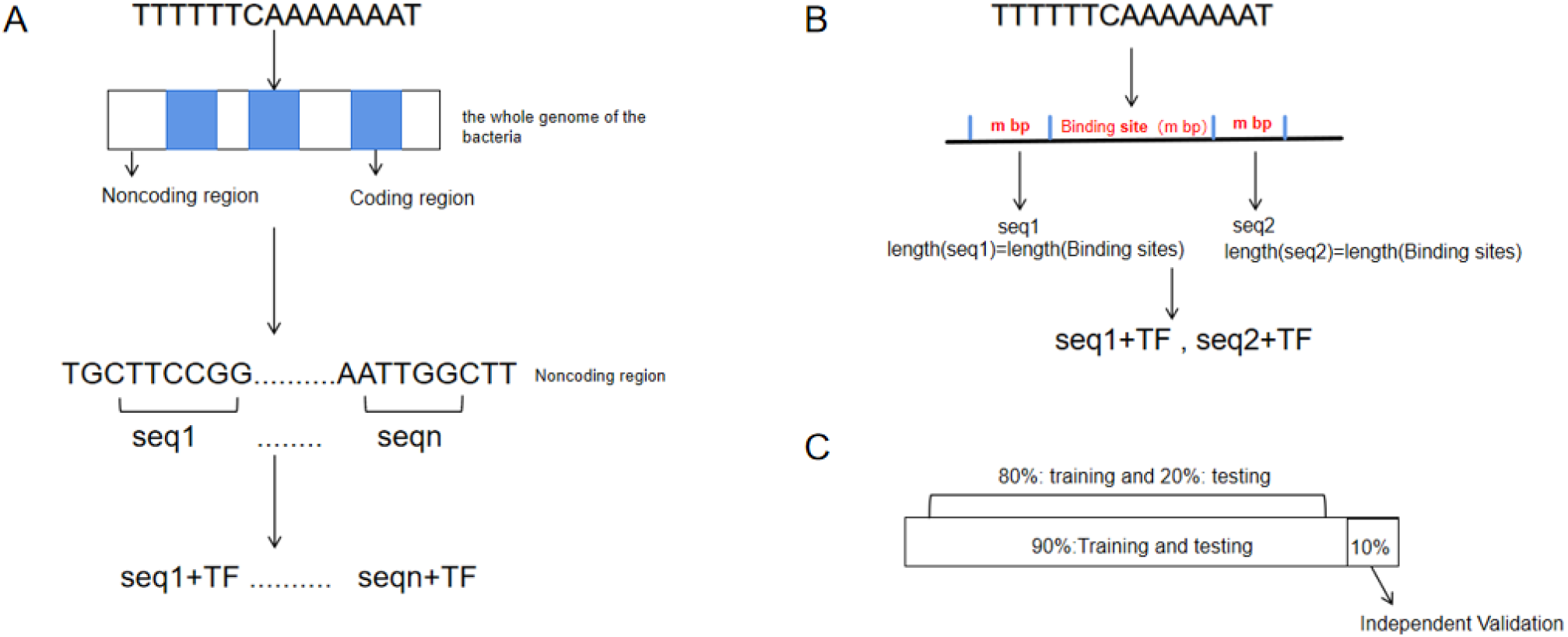
Negative sample extraction methods. A. Flowchart of the RS. B. Flowchart of the EE. C. Schematic representation of the datasets.

**Figure2.**
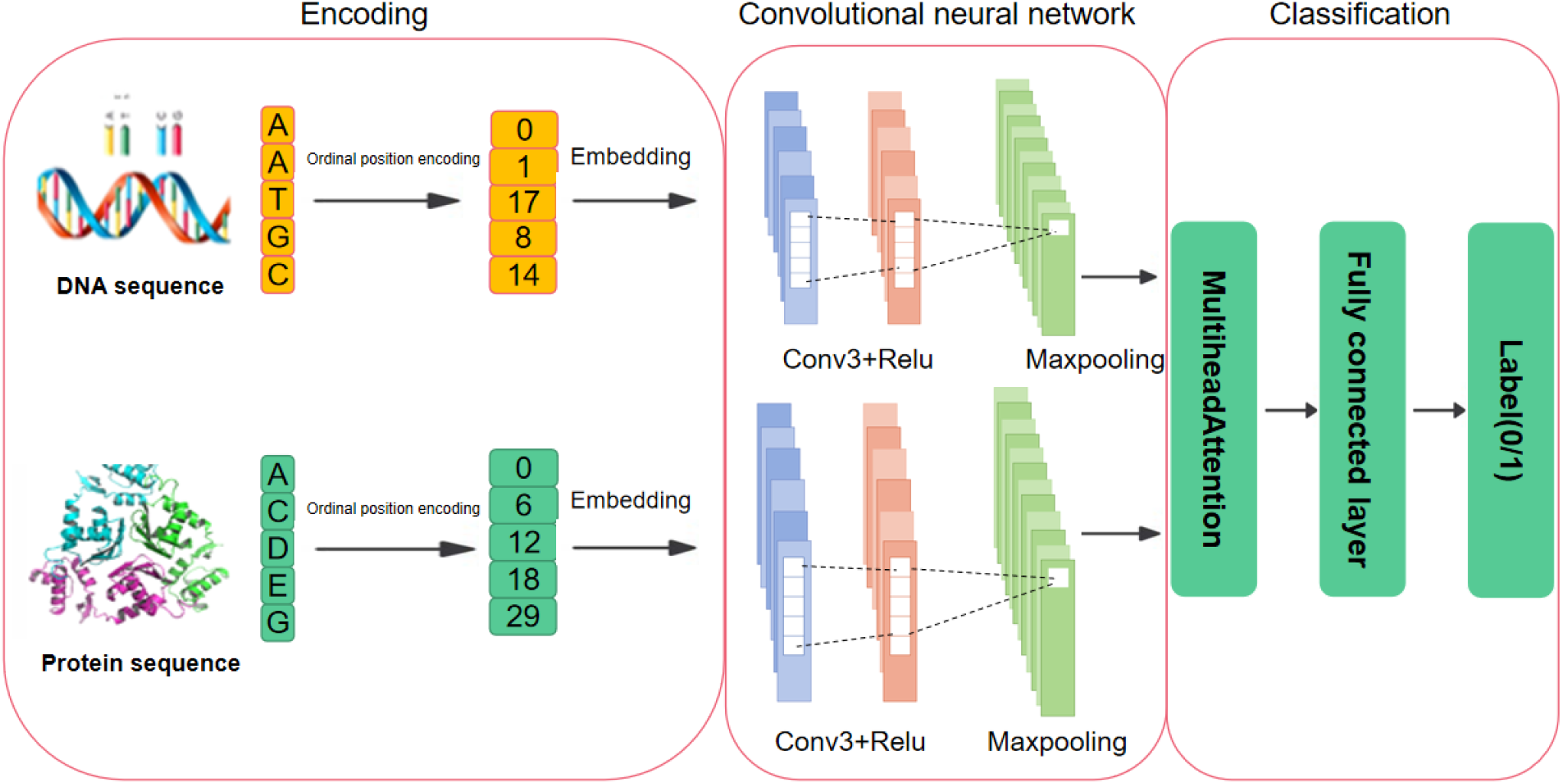
Flowchart of bacteria transcription factors and its binding sites BTFBS model based on convolutional neural network and MultiheadAttention.

The second method (EE) is based on the experimental experience: the adjacent positions of the binding sites are not combined with their corresponding transcription factors. CollecTF provides the location information of binding sites. The DNA sequences are obtained by intercepting the adjacent positions of the binding sites, resulting in sequences of equal length to the binding sites themselves. Consequently, 9112 non-redundant sequences are obtained, with 57% of the data randomly selected as negative samples, in order to match the number of positive samples. Additionally, 10% of the sequences are randomly selected as the independent datasets, while the remaining 90% are utilized as the training and test datasets. Figure 1B illustrates the process of EE.

The positive and negative samples are merged to form the total datasets, which is then divided into an 80% training set and a 20% test set. Five-fold cross-validation experiments are conducted on the training set (Figure 1C).

### Model construction

In this work, we utilize two negative sample sampling methods, and use convolutional neural networks as well as MultiheadAttention to construct a predictive model, BTFBS. The construction of BTFBS for bacteria is illustrated in Figure2. The model has four main parts. The first part is the encoding module, which converts the sequence into dense vectors of fixed size. The second part is the convolutional neural network module, which extracts features. The third part is the MultiheadAttention module, which captures connections and differences in the sequences. The fourth part is the classification module, which gives the classification results. The pipeline of BTFBS is shown in Algorithm 1.

#### Algorithm 1

The pipeline of BTFBS

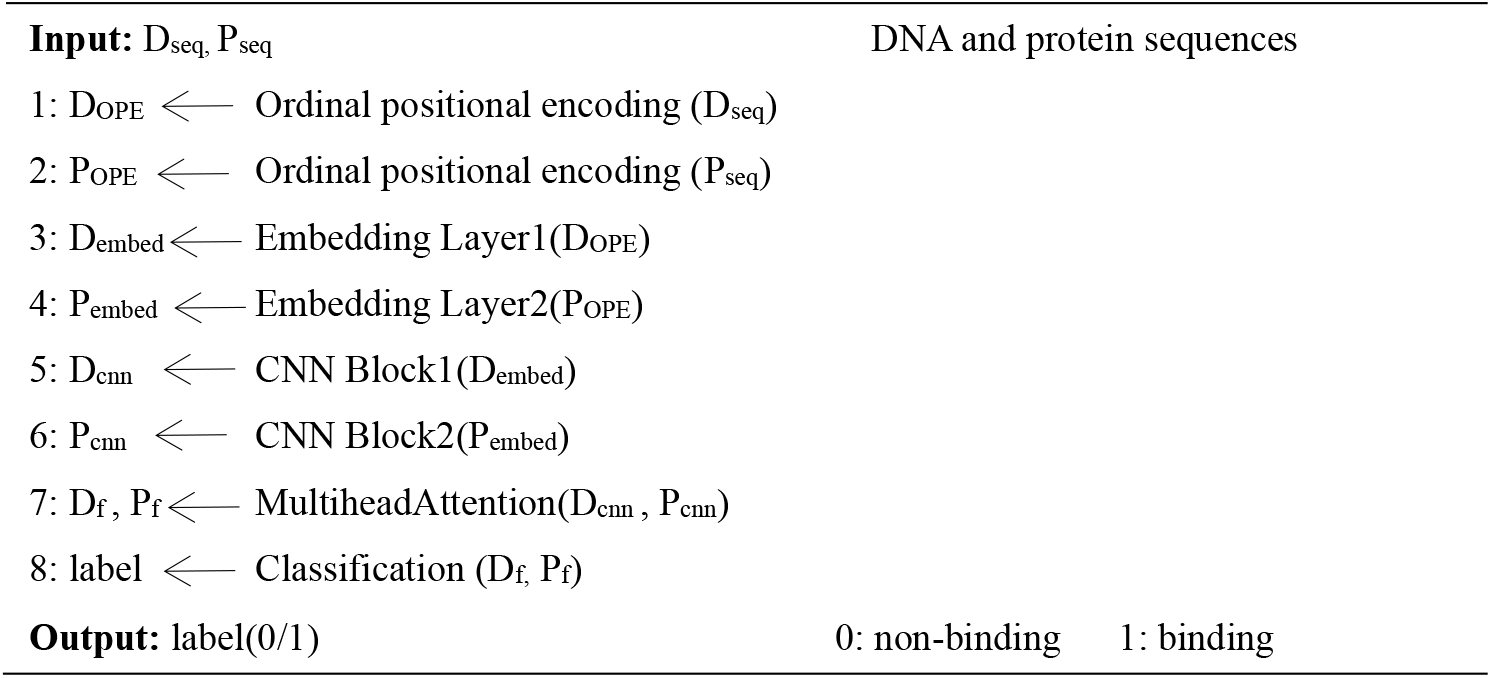

#### Encoding

Before embedding, we first carry out the ordinal positional encoding on the sequences. It is known from the literature that ordinal positional encoding [10] can improve the accuracy of CNN models and cross-validation, so that each sequence is represented as integers. In order to ensure the consistency of sequence length, the maximum length of a DNA sequence is set to 100 and the maximum length of a protein sequence is set to 600. Sequences shorter than the maximum length are padded with zeros to match the fixed length. Then the embedding layer receives these fixed length vectors and converts them into dense continuous feature vectors with a fixed size. As the first layer of embedding training, the weight matrix is constantly updated during the training process [14], and eventually an embedding matrix is learned to represent the sequence.

#### CNN

The previously learned embedding matrix is successively fed into convolution layer with three convolution kernels, and the rectified linear unit (ReLU) is used as the activation function. Finally, the obtained eigenvectors are processed by the maximum pooling layer.

#### MultiheadAttention

Similar to that described by Bian et al. [15], the architecture of MultiheadAttention is described as follows. The DNA attention component first passes a DNA feature map D_cnn_ through a linear layer to compute a DNA query vector Q^i^_D_ , a DNA key vector K^i^_D_ and a DNA value vector V^i^_D_ . The queries, keys and values for the DNA are given by

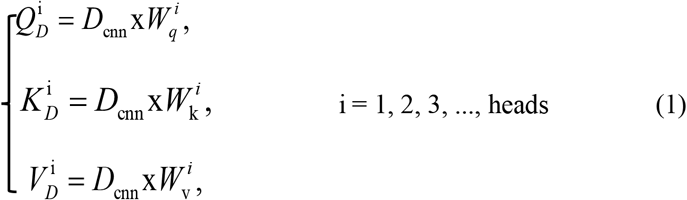

where W^i^_q_, W^i^_k_, W^i^_v_ are the weight matrices of three linear layers, and heads is the number of attention heads.

The protein attention component follows a similar process to the DNA attention component, and both attention components shares these three linear layers. The protein attention component first delivers a protein feature map P_cnn_ through a linear layer to compute a protein query vector Q^i^_P_ , a protein key vector K^i^_P_ and a protein value vector V^i^_P_ . The queries, keys and values for the protein are given by

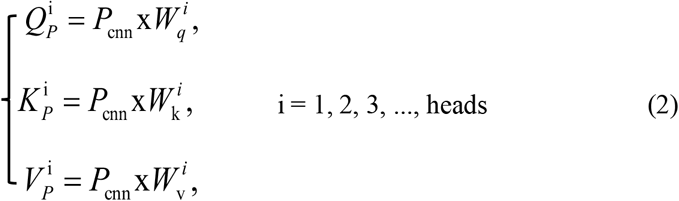

where W^i^_q_ , W^i^_k_ , W^i^_v_ are the weight matrices of three linear layers, and heads is the number of attention heads. The attention matrix for DNAs and proteins is calculated as follows by using the softmax function:

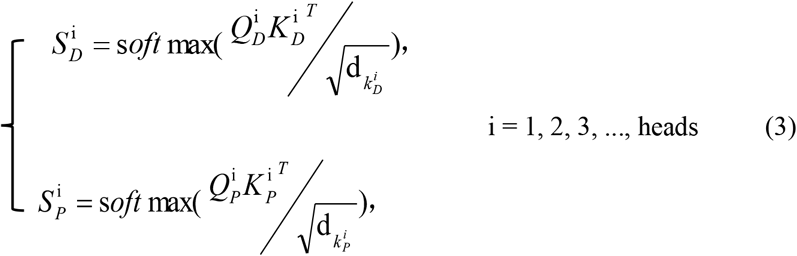

where 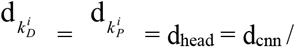 heads is the dimension of queries and keys.

The DNA attention matrix for each head is multiplied by the DNA value matrix for each head to obtain the attended DNA feature map for each head. Then the attended DNA feature maps of all attention heads are concatenated along the channel dimension and fed into a linear layer to obtain the final attended DNA feature map Z_DC_:

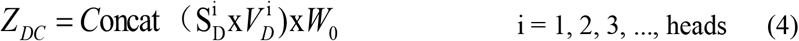

where W_0_ is weight matrix.

The protein attention matrix S^i^_P_ and protein value matrix V^i^_P_ are operated in the same way to get the final protein feature map Z_PC_:

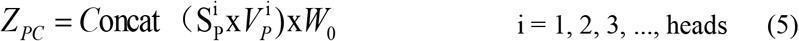

where W_0_ is weight matrix.

Finally, in order to obtain the final feature maps, the attended feature maps are combined with the original feature maps:

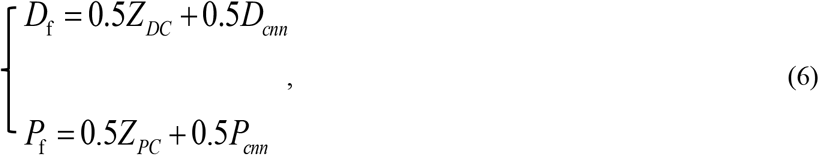

where D_f_ denotes the DNA feature map and P_f_ denotes the protein feature map.

#### Classification

We use the fully connected layer as the classification module, and concatenate the feature vectors obtained by convolutional neural network into the classification module. The classification module consists of dropout layer, fully connected layer and ReLU activation function. The dropout layer [15] is used to avoid overfitting during training.

### Evaluation metrics

In order to measure the performance of the models, we calculate the accuracy (ACC), sensitivity, specificity, F1-score, and Matthews correlation coefficient (MCC), which are defined below [16]:

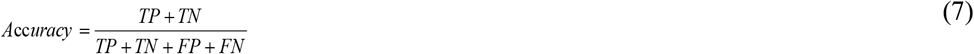

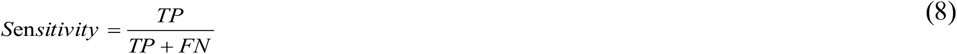

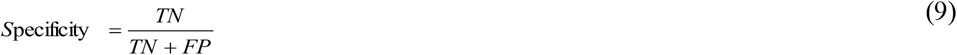

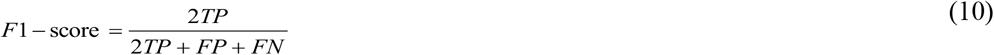

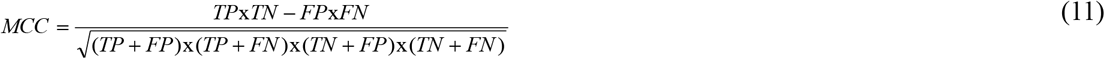

where TP is true positive, TN is true negative, FP is false positive, and FN is false negative.

## Results and discussion

### Comparison of models based on the two negative sample sampling methods

This paper employs two distinct negative sampling methods: RS and EE. As shown in Table1 and Table2, the results on test dataset for RS are found to be ACC=0.91446, Sensitivity=0.89746, Specificity=0.93134, MCC=0.82946, F1-score=0.91264, while the results of EE are that ACC=0.87868, Sensitivity=0.89354, Specificity=0.86394, MCC=0.75796, F1-score=0.87996. As seen in Figure 3A and Figure 3B, the performance of RS is better than that of EE, and the accuracies of independent validation and testing are similar, indicating no bias for the prediction models. But for the merged independent datasets (RS+EE), EE shows better prediction accuracy than RS, indicating the prediction model trained by EE has better generalization ability than RS.

**Figure3.**
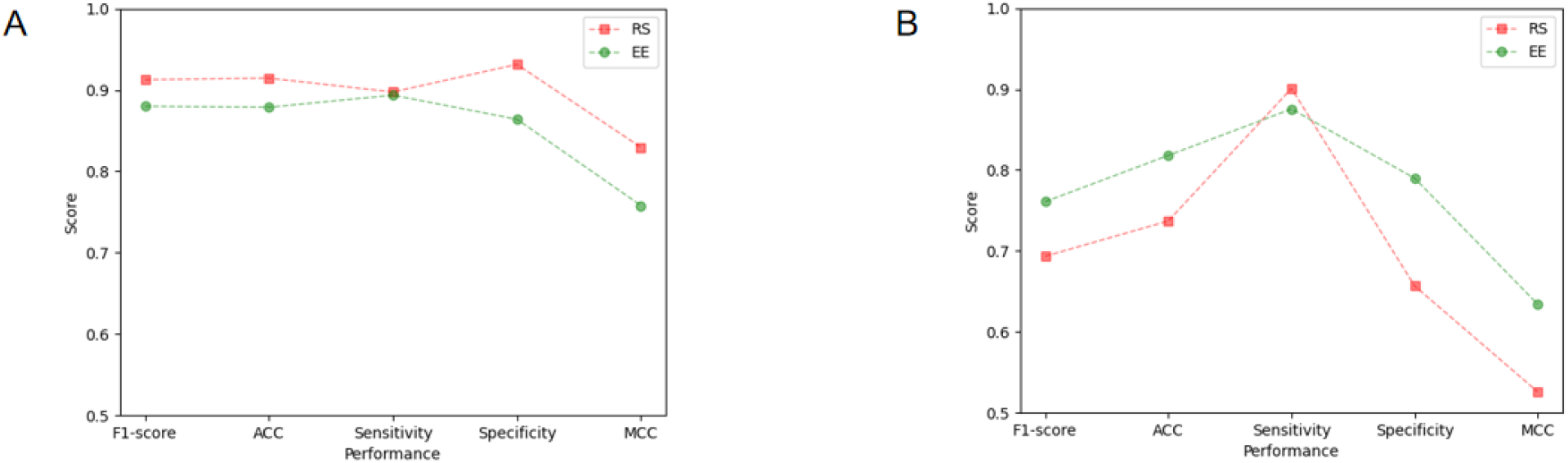
A. Comparison of models based on the two negative sample sampling methods on test data set. B. Comparison of models based on the two negative sample sampling methods on merged independent datasets (RS+EE).

**Table1.**
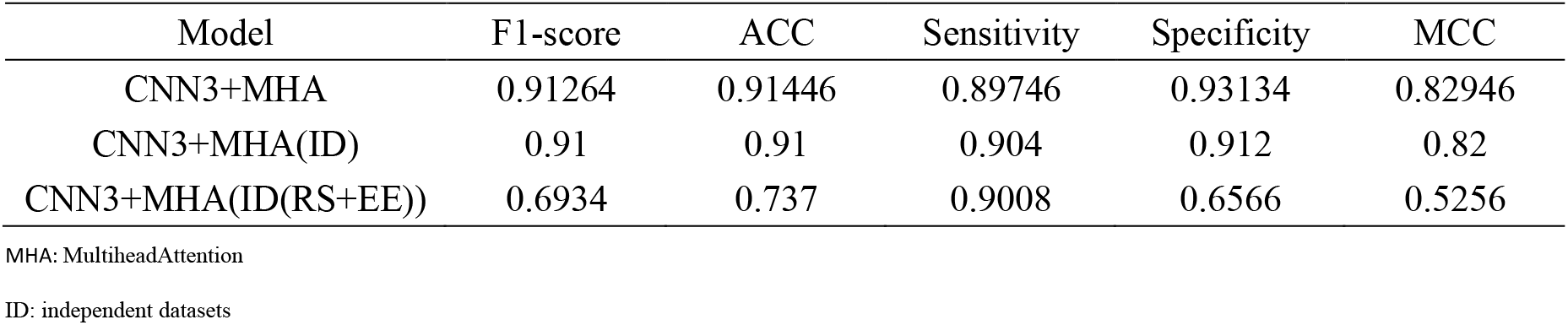
The performance of the model on the various datasets (RS)

| Model | F1-score | ACC | Sensitivity | Specificity | MCC |
| --- | --- | --- | --- | --- | --- |
| CNN3+MHA | 0.91264 | 0.91446 | 0.89746 | 0.93134 | 0.82946 |
| CNN3+MHA(ID) | 0.91 | 0.91 | 0.904 | 0.912 | 0.82 |
| CNN3+MHA(ID(RS+EE)) | 0.6934 | 0.737 | 0.9008 | 0.6566 | 0.5256 |

MHA: MultiheadAttention
ID: independent datasets

**Table2.** The performance of the model on the various datasets (EE)

| Model | F1-score | ACC | Sensitivity | Specificity | MCC |
| --- | --- | --- | --- | --- | --- |
| CNN3+MHA | 0.87996 | 0.87868 | 0.89354 | 0.86394 | 0.75796 |
| CNN3+MHA(ID) | 0.876 | 0.876 | 0.888 | 0.864 | 0.76 |
| CNN3+MHA(ID(RS+EE)) | 0.7608 | 0.818 | 0.8754 | 0.7898 | 0.6338 |

MHA: MultiheadAttention
ID: independent datasets

#### Ordinal positional encoding can improve the performance of the model

Ordinal positional encoding is a coding method that adds positional information [10]. To test whether this coding method improves the performance of the model, a comparative test is conducted. The results of RS and EE are shown in Figure 4A and Figure 4B, respectively. It is clear that a model with ordinal positional encoding performs better than the one without it.

**Figure4.**
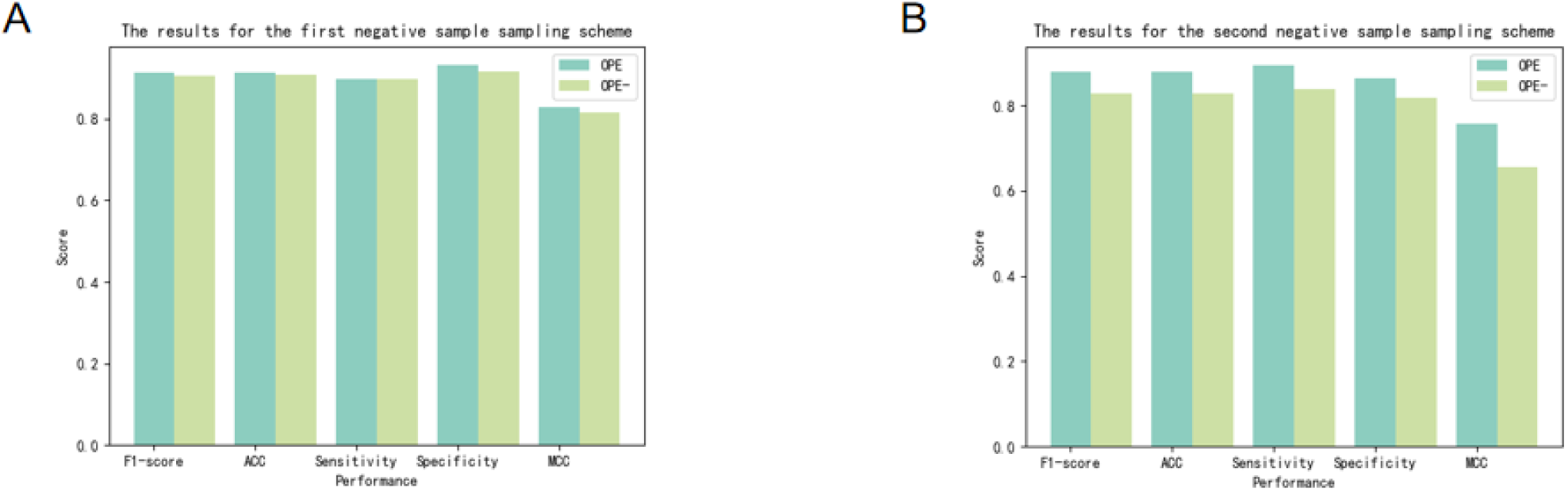
Results on test sets with and without the ordinal positional encoding. A. Results of RS. B. Results of EE.

#### Comparison with some other methods

The performance of BTFBS is compared with several usual methods for predicting protein-DNA-binding sites [17], including DeepBind [2], Dilated [3], KEGRU [4], DanQ [5], and MAResNet [6]. DeepBind, Dilated, DanQ as well as KEGRU were implemented based on the deepRAM [9] framework. Since our model has two input parts, to ensure fairness, we modify BTFBS to take only one input of DNA sequence for comparison with these models. Bacterial binding sites are used as positive samples (while negative samples are obtained through deepRAM), and they are randomly divided into a training set comprising 80% and a test set comprising 20%. The aforementioned data are then used to train DeepBind, Dilated, DanQ, KEGRU and MAResNet, respectively. Following the completion of three-fold cross-validation on the training set and subsequent evaluation of the models’ performance on the test dataset, the two metrics for the different models on the test datset are presented in Figure 5A, and Table 3 lists additional performance metrics for these models. The accuracy and AUC of BTFBS are 0.8065 and 0.8854, representing an improvement of 0.86% in accuracy and 3.72% in AUC compared to the second highest performing model.

**Table3.** The performance of BTFBS and other methods on the test set (shuffle)

| Model | Accuracy | Sensitivity | Specificity | MCC | F1 | AUC |
| --- | --- | --- | --- | --- | --- | --- |
| DeepBind | 0.7272 | 0.7560 | 0.7068 | 0.4563 | 0.6966 | 0.8105 |
| Dilated | 0.7878 | 0.7884 | 0.7872 | 0.5752 | 0.7961 | 0.8482 |
| DanQ | 0.7878 | **0.8163** | 0.7600 | 0.5770 | 0.7920 | 0.8207 |
| KEGRU | 0.7979 | 0.7454 | **0.8636** | 0.6060 | **0.8039** | 0.8176 |
| MAResNet | 0.7337 | 0.6993 | 0.7575 | 0.4572 | 0.7407 | 0.8368 |
| BTFBS | **0.8065** | 0.8004 | 0.8124 | **0.6131** | 0.8012 | **0.8854** |

**Figure5.**
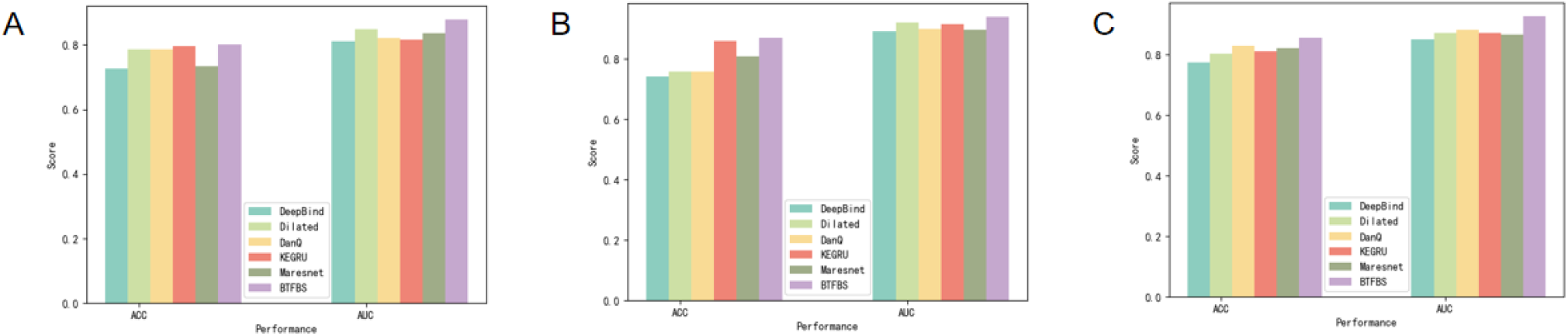
The results of 3-fold cross-validation of BTFBS and some other existing methods. A. The negative sample comes from shuffling the positive sample. B. The negative sample comes from RS. C. The negative sample comes from EE.

Subsequently, the negative samples are replaced with data from RS and EE, respectively, to ascertain whether these two negative sample schemes are more effective than the shuffling method. The training of the model is conducted in accordance with the same procedure as that employed for the shuffling method. Figure 5B depicts the two metrics (RS) for the various models on the test dataset, while Table 4 enumerates supplementary performance metrics for these models. Figure 5C illustrates the two metrics (EE) for the disparate models on the test set, with Table 5 providing additional performance metrics for these models. We can see from the above results: (1) Our model outperforms other models in all three negative sample sampling methods. (2) When using negative sample data in RS or EE, each model performs better than using the shuffling method. A possible reason is that the data obtained by the shuffling method may be not pieces of the whole genome sequences of the corresponding bacteria.

**Table4.**
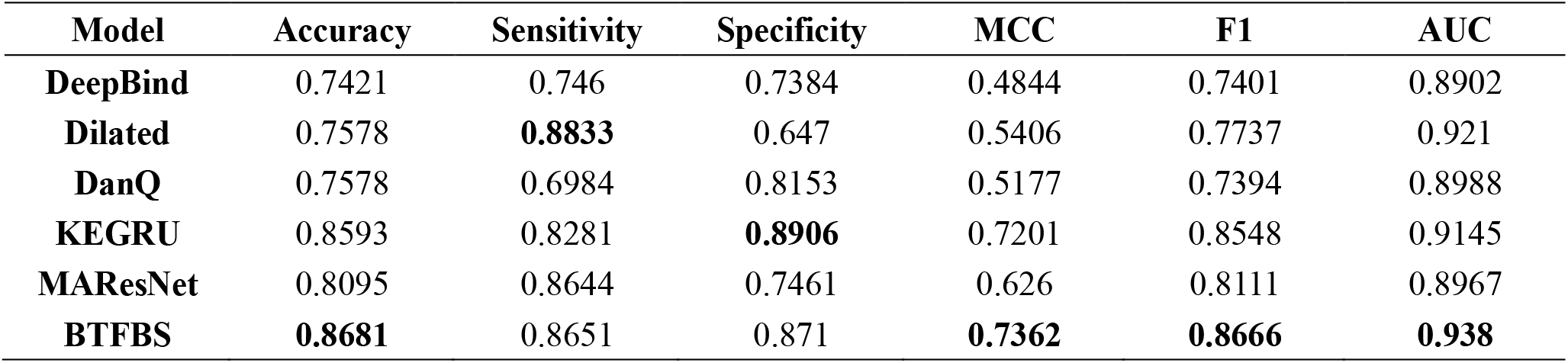
The performance of BTFBS and other methods on the test set (RS)

| Model | Accuracy | Sensitivity | Specificity | MCC | F1 | AUC |
| --- | --- | --- | --- | --- | --- | --- |
| DeepBind | 0.7421 | 0.746 | 0.7384 | 0.4844 | 0.7401 | 0.8902 |
| Dilated | 0.7578 | **0.8833** | 0.647 | 0.5406 | 0.7737 | 0.921 |
| DanQ | 0.7578 | 0.6984 | 0.8153 | 0.5177 | 0.7394 | 0.8988 |
| KEGRU | 0.8593 | 0.8281 | **0.8906** | 0.7201 | 0.8548 | 0.9145 |
| MAResNet | 0.8095 | 0.8644 | 0.7461 | 0.626 | 0.8111 | 0.8967 |
| BTFBS | **0.8681** | 0.8651 | 0.871 | **0.7362** | **0.8666** | **0.938** |

**Table5.**
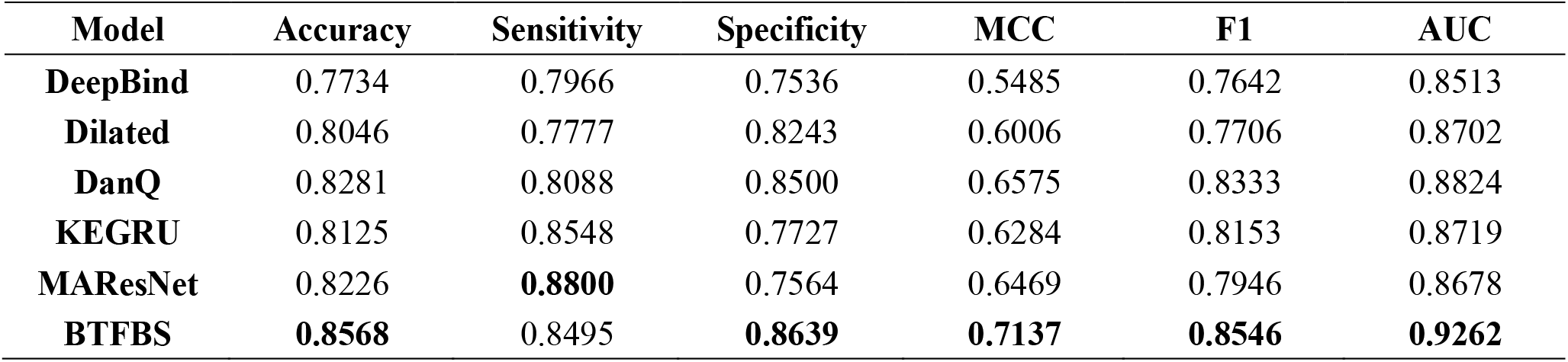
The performance of BTFBS and other methods on the test set (EE)

| Model | Accuracy | Sensitivity | Specificity | MCC | F1 | AUC |
| --- | --- | --- | --- | --- | --- | --- |
| DeepBind | 0.7734 | 0.7966 | 0.7536 | 0.5485 | 0.7642 | 0.8513 |
| Dilated | 0.8046 | 0.7777 | 0.8243 | 0.6006 | 0.7706 | 0.8702 |
| DanQ | 0.8281 | 0.8088 | 0.8500 | 0.6575 | 0.8333 | 0.8824 |
| KEGRU | 0.8125 | 0.8548 | 0.7727 | 0.6284 | 0.8153 | 0.8719 |
| MAResNet | 0.8226 | **0.8800** | 0.7564 | 0.6469 | 0.7946 | 0.8678 |
| BTFBS | **0.8568** | 0.8495 | **0.8639** | **0.7137** | **0.8546** | **0.9262** |

**Table6.**
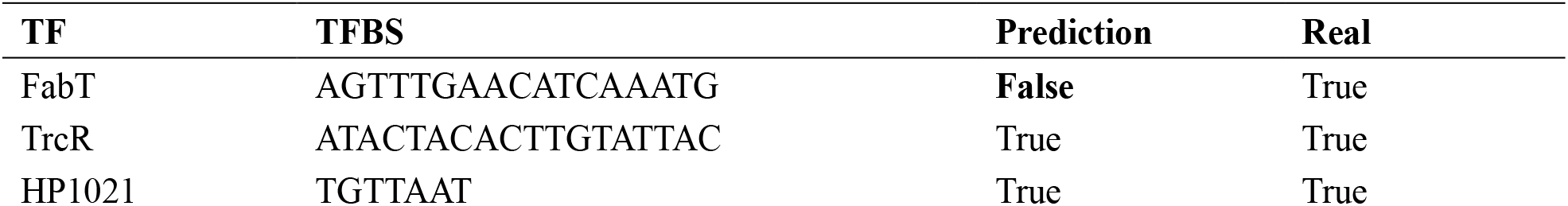
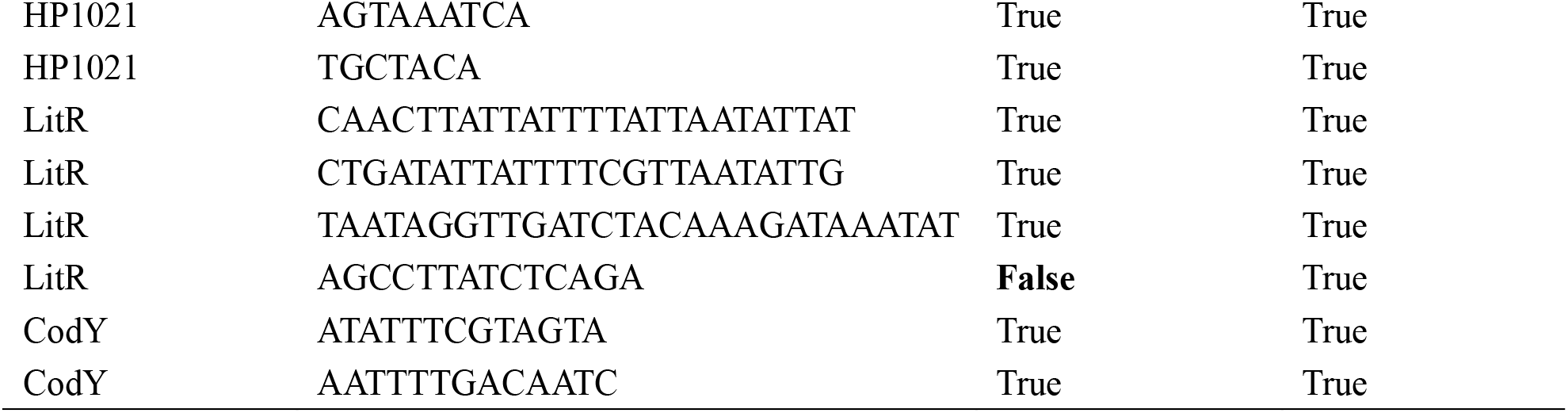
The prediction results of RS.

### Case Study

In order to demonstrate the validity and generalization ability of our proposed method, a search of the literature from 2022 to 2024 on the Web of Science is conducted using the keywords “bacteria”, “transcription factor”, “binding sites”. A total of 11 pieces of data are retrieved, which are able to identify protein sequences of transcription factors and DNA sequences of binding sites. These include FabT [18], TrcR [19], HP1021 [20], LitR [21] and CodY [22]. In RS, two prediction errors are identified in the positive samples. The EE has two instances of incorrect prediction for positive samples.

### BTFBS provides an experimental guide

While experimental methods for obtaining transcription factor binding data are highly accurate, these methods are also time-consuming, laborious, and costly. Our model BTFBS can guide experiments and help researchers to narrow down the scope of their work.

WOAH Reference Lab for Swine Streptococcosis, College of Veterinary Medicine, Nanjing Agricultural University provides two 100 bp DNA fragments that have been demonstrated to bind to the transcription factors XtrSs, ArgR, respectively. In order to select an appropriate sliding window length, we have counted the lengths of all binding sites in CollecTF (Figure 6A) as well as those of experimentally studied Streptococcus (Figure 6B). Figure 6 illustrates that the majority of binding sites are distributed between 6 and 30 bp in length, whereas the binding sites of Streptococcus are predominantly distributed between 6 and 25 bp. Subsequently, the aforementioned DNA sequences are intercepted using a sliding window of length 25 and a step size of 1, respectively. For XtrSs, the RS-predicted fragment containing the binding site scored 0.99009, ranking third out of 76 DNA fragments (Table 8), while the EE-predicted fragment scored 0.9874, ranking second out of 76 DNA fragments (Table 9). For ArgR, the RS-predicted fragment containing the binding site scored 0.9994, ranking second out of 76 DNA fragments (Table 10), while the EE-predicted fragment scored 0.9172, ranking twelfth out of 76 DNA fragments (Table 11). Specific binding site information is available in [23].

**Table7.**
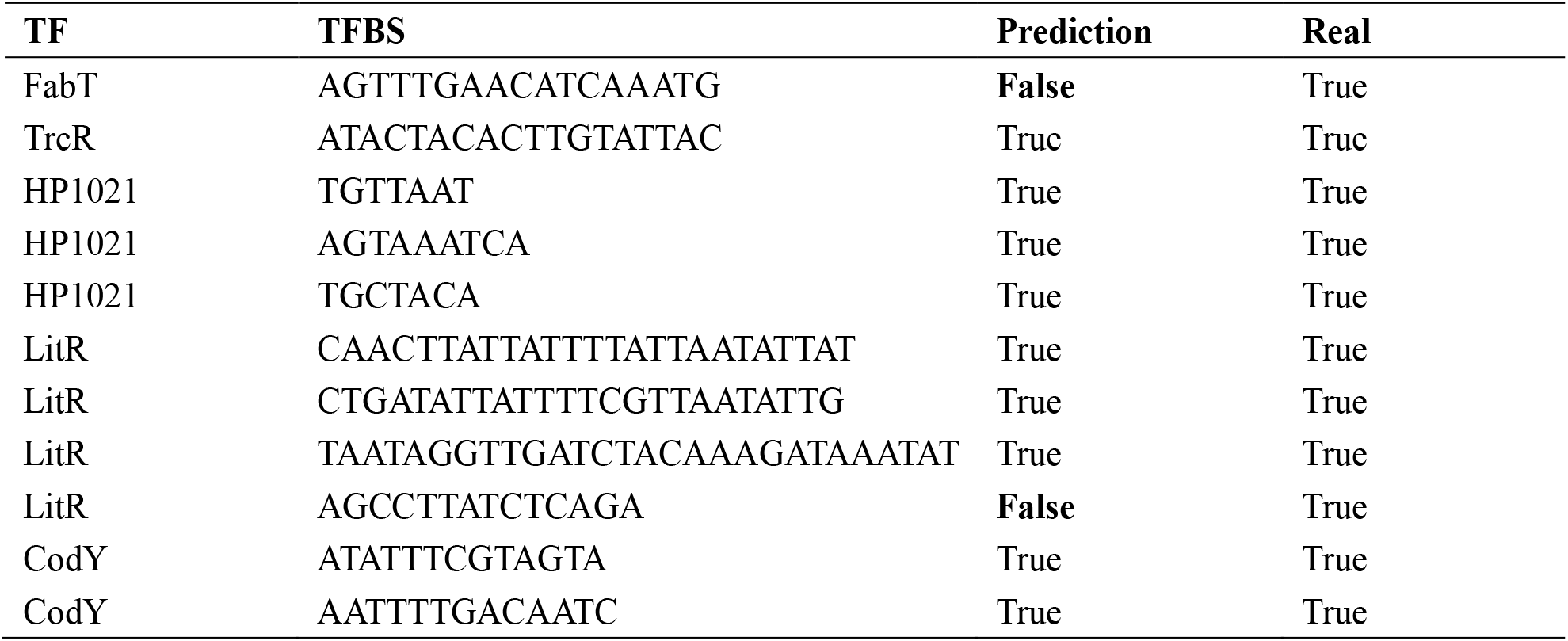
The prediction results of EE.

**Table8.**
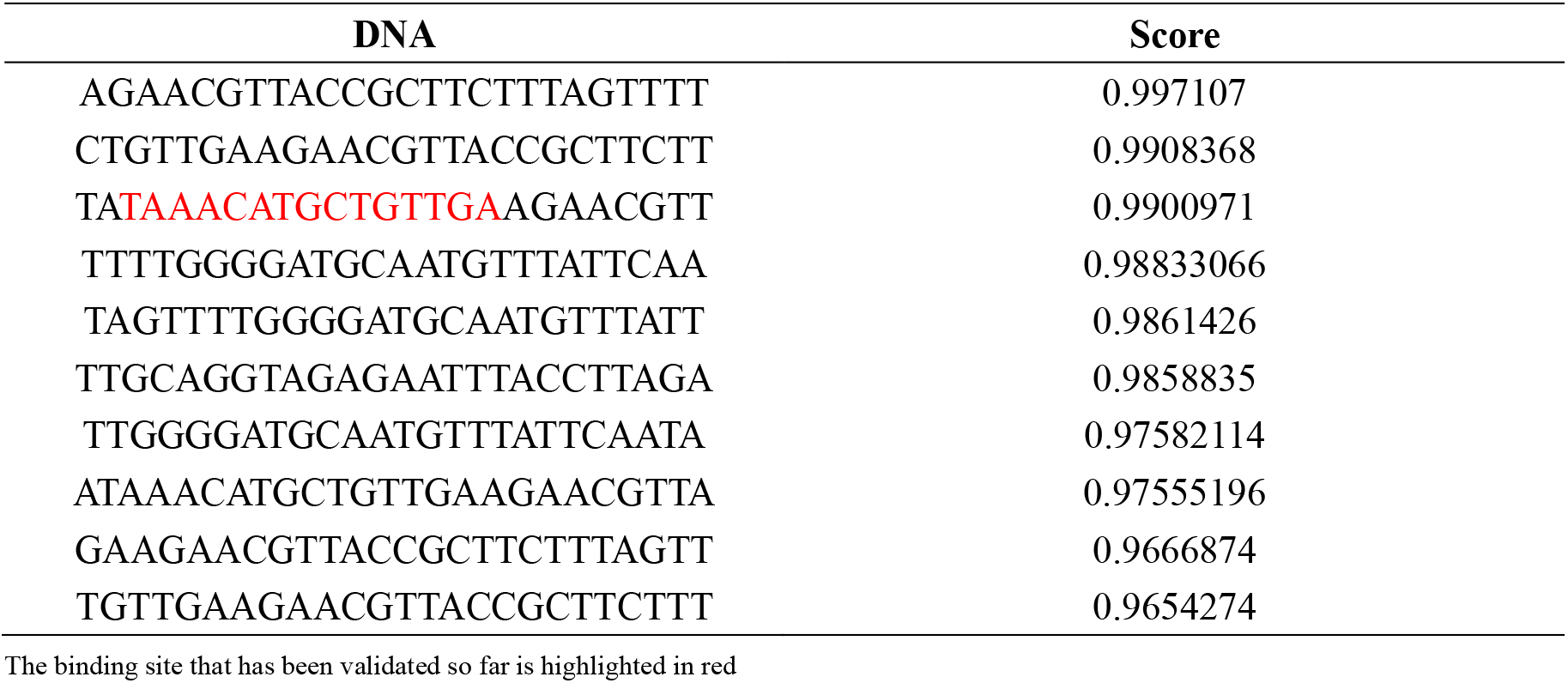
DNA prediction score (XtrSs) by using RS.

**Table9.**
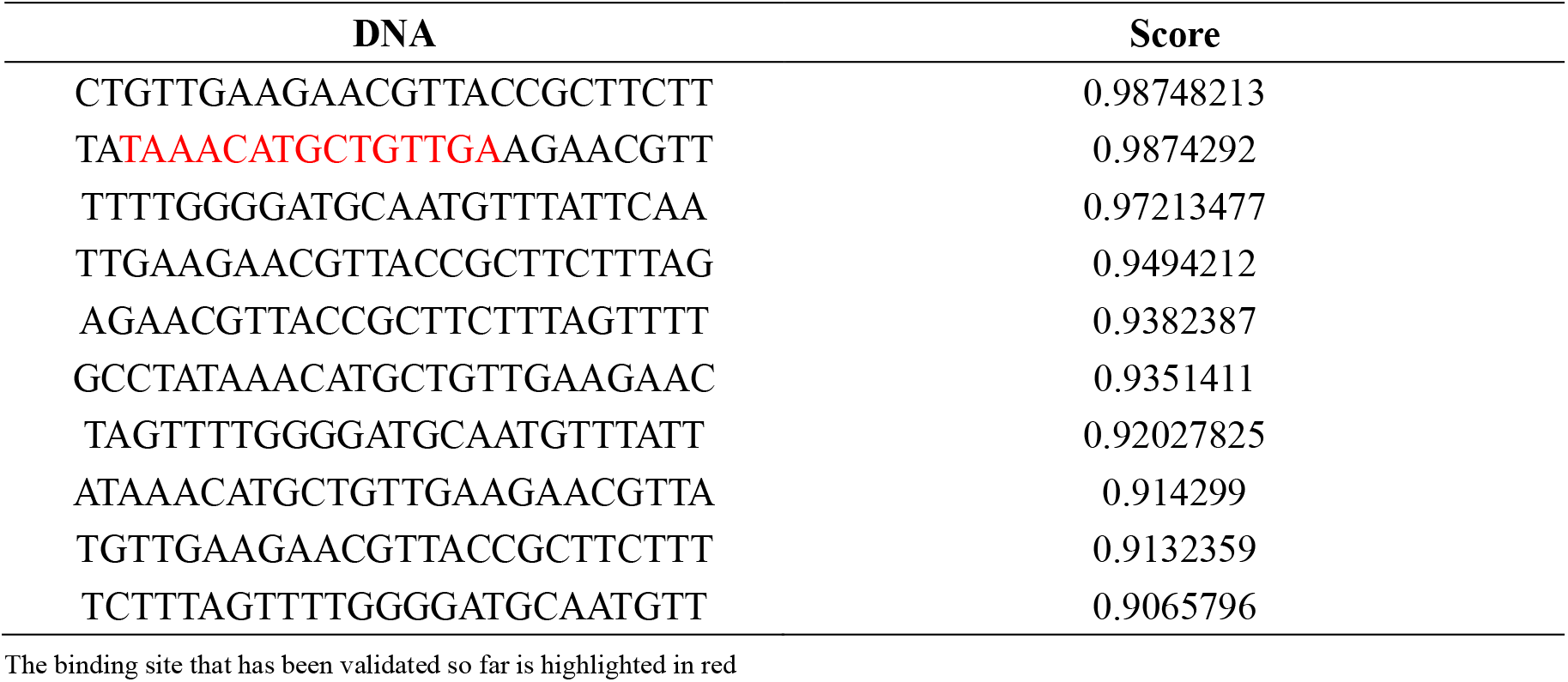
DNA prediction score (XtrSs) by using EE.

**Table10.**
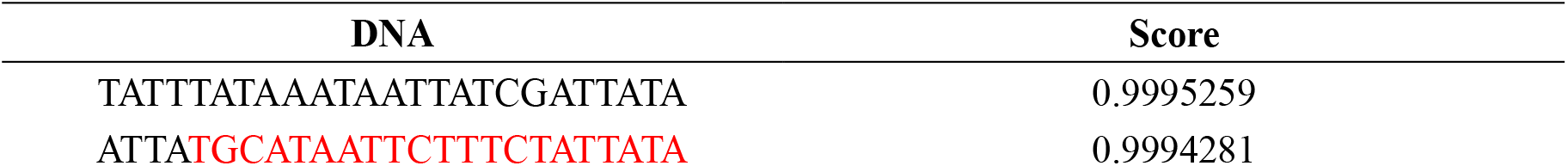
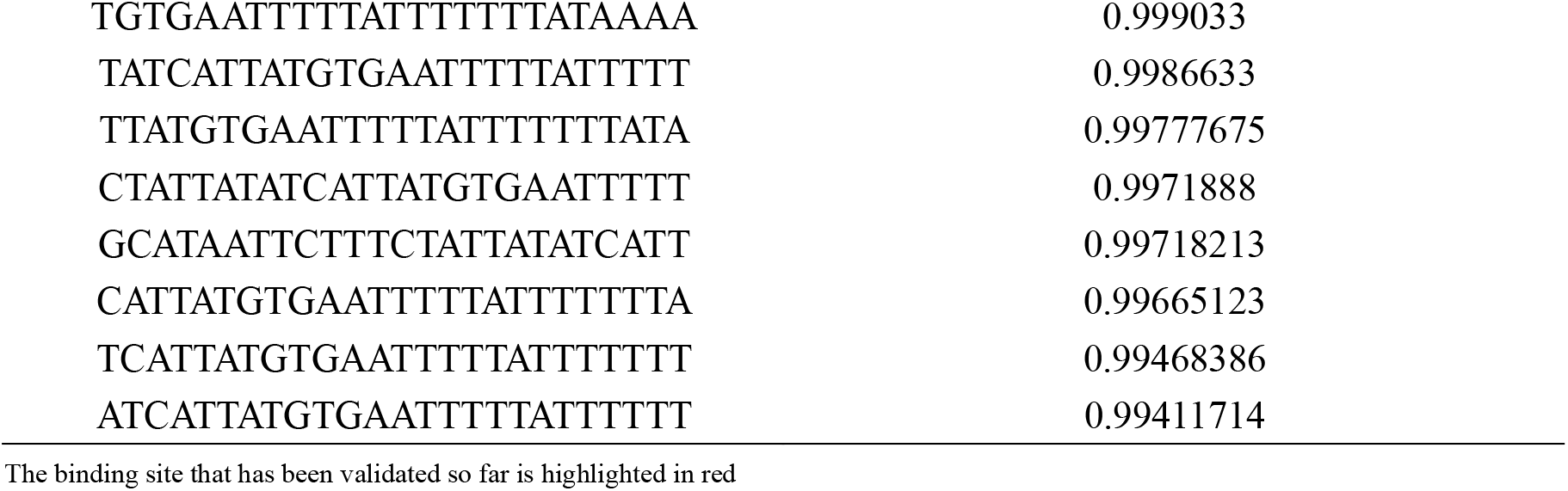
DNA prediction score (ArgR) by using RS.

**Table11.**
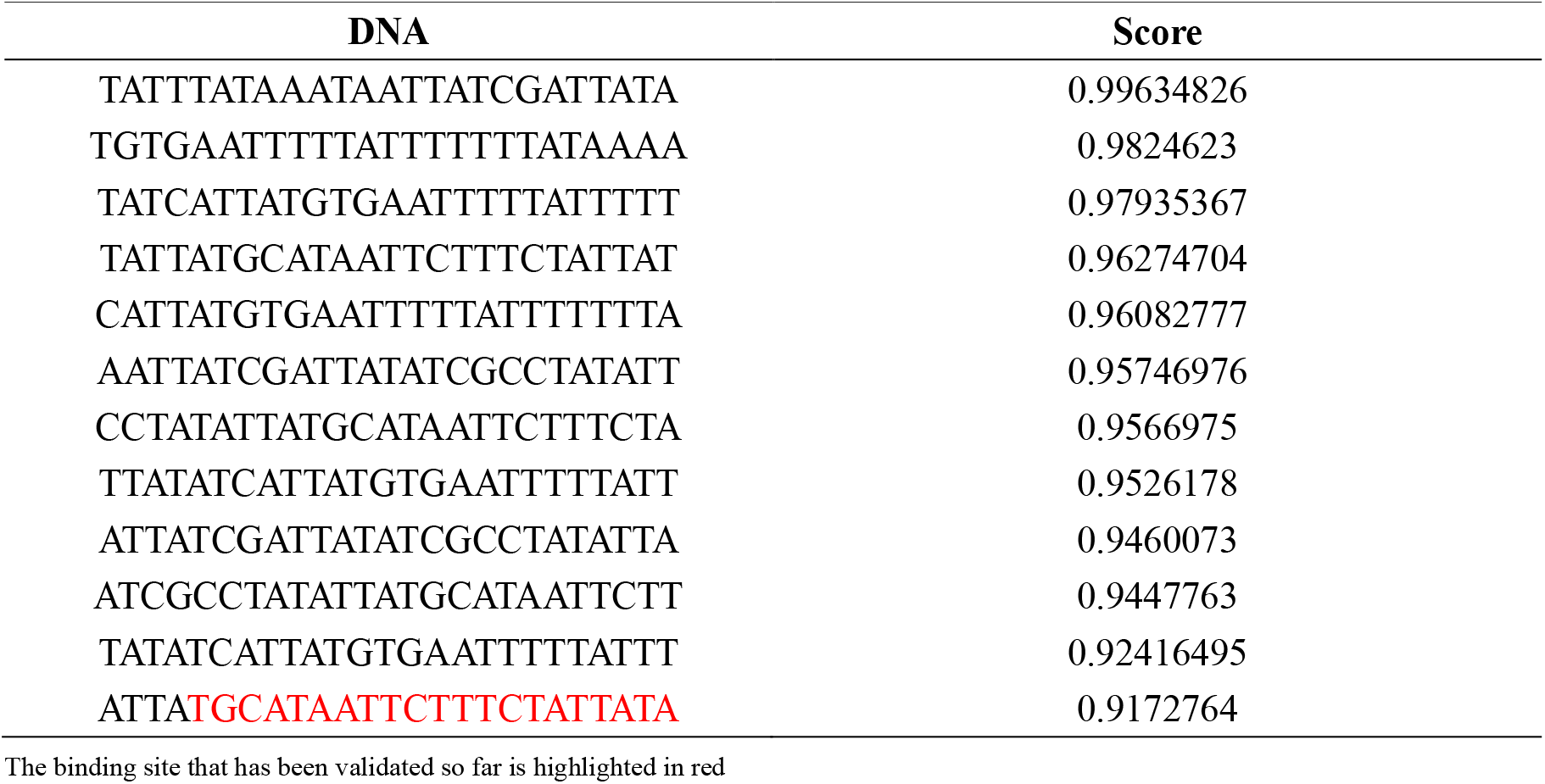
DNA prediction score (ArgR) by using EE.

**Figure6.**
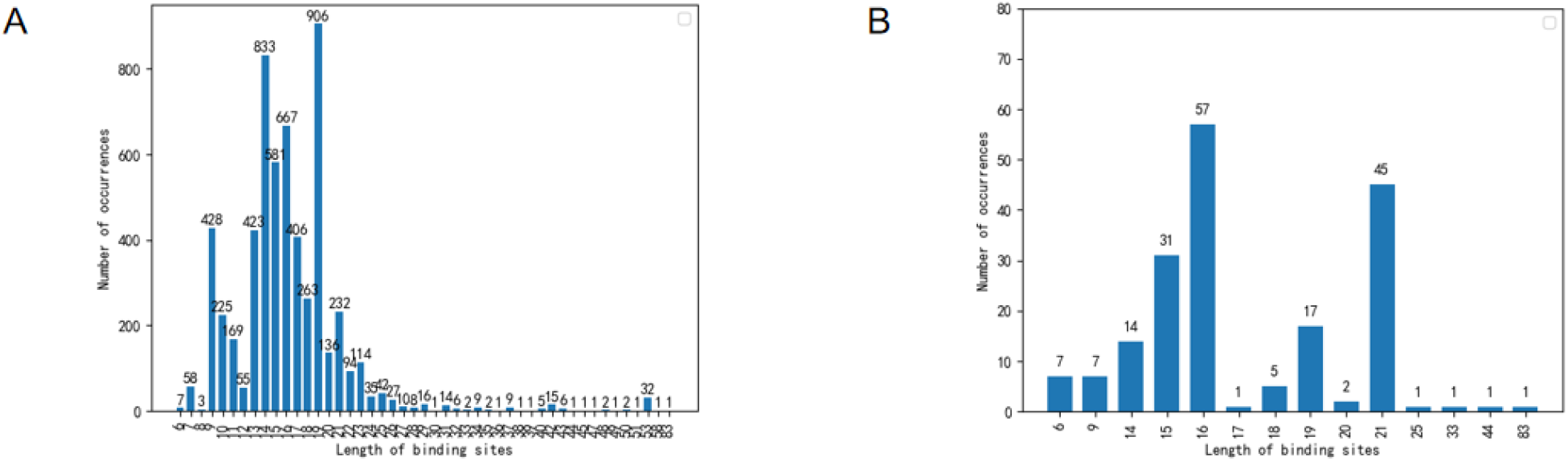
Binding sites length distribution map. A. Distribution of all binding sites lengths. B. Streptococcus binding sites length distribution.

## Discussion

In order to predict whether bacterial transcription factors and binding sites bind or not, we propose a model BTFBS, which takes protein sequences of transcription factors and nucleotide sequences of binding sites as inputs. BTFBS, based on deep learning, can automatically extract features from training data and capture the internal relationship between protein and nucleotide sequences. The experimental results show that ordinal positional encoding can improve the performance of the model by adding positional information. Two negative sample sampling methods are adopted, and it is found that the choice of negative samples has an impact on the model’s performance. The performance of RS is better than that of EE. But the model trained by EE has better generalization ability than RS. One possible reason is that in RS, there is a difference between the randomly truncated DNA fragments and the binding sites, making it easier for the model to learn this difference, whereas in EE, the DNA fragments are truncated from the adjacent positions of the binding sites, which have some connection to the binding sites, making it difficult for the model to distinguish them.

## Conclusion

We put forward a model named BTFBS, which has two inputs: protein sequences of transcription factors and nucleotide sequences of binding sites. Our work mainly includes the following four points: (1) We provide a deep learning model with two inputs that can capture links between transcription factors and binding sites. (2) Ordinal positional encoding is introduced to add positional information, which can improve the performance of the model.

(3) We use two negative sample sampling methods. For bacteria, the model performance of RS and EE compared to the shuffling method indicates that the negative samples should be come from the whole genome sequences of the corresponding bacteria. (4) Our proposed method is superior to some usual methods for predicting protein-DNA-binding sites.

Although the results show that BTFBS has good performance, there is still room for further improvement: (1) Contrastive learning can maximize the agreement (disagreement) between similar (dissimilar) samples [24]. Later, we will consider introducing contrast learning to further optimize the model performance. (2) We might consider studying only one type of bacteria, such as Escherichia coli. (3) Oliveira Monteiro et al. trained the deep learning model PredicTF by constructing a bacterial transcription factors database. PredicTF can realize the prediction of bacterial transcription factors [5]. We plan to use PredicTF to predict the likely transcription factors of a bacteria and then split its whole genome sequence using sliding windows. We could use our model to make prediction scores for predicted transcription factors and DNA fragments.

## Abbreviations

TF: Transcription factor
CNN: Convolutional neural network
BTFBS: Bacterial Transcription Factor and Binding Site
ReLU: Rectified linear unit
ACC: Accuracy
MCC: Matthews correlation coefficient

## Acknowledgements

Not applicable.

## Author contributions

T Y and GJ L developed the concept of the study. T Y、GJ L、S L、YY C and BB J did the analyses. S L、XQ L、R Z、Y Z and WQ Z collated data. BB J performed the experiments, prepared figures and tables, drafted the manuscript. GJ L and TY critically revised the manuscript and finalized the submission. All co-authors read and approved the final manuscript.

## Availability of data and materials

Codes and Datasets are available at https://github.com/Vceternal/BTFBS

## Declarations

### Ethics approval and consent to participate

Not applicable.

### Consent for publication

Not applicable.

